# Enhanced gene expression plasticity as a mechanism of adaptation to a variable environment in a reef-building coral

**DOI:** 10.1101/059667

**Authors:** C.D. Kenkel, M.V. Matz

## Abstract

Local adaptation is ubiquitous^1^, but the molecular mechanisms giving rise to this ecological phenomenon remain largely unknown^2^. A year-long reciprocal transplant of mustard hill coral (*Porites astreoides*) between a highly environmentally variable inshore habitat and more stable offshore habitat^3^ demonstrated that both inshore and offshore populations exhibit elevated growth, protein and lipid content in their home reef environment, indicative of local adaptation ^4^ Here, we characterized the genomic basis of this adaptation in both coral hosts and their intracellular algal symbionts (*Symbiodinium* sp.) using genome-wide gene expression profiling^5,6^ and gene coexpression network analysis^7^. Inshore and offshore coral populations differ primarily in their capacity for gene expression plasticity: upon transplantation to a novel environment inshore corals were able to match expression profiles of the local population significantly better than offshore corals. Furthermore, elevated plasticity in expression of environmental stress response (ESR) genes was adaptive in the inshore environment: it correlated with the least susceptibility to a natural summer bleaching event, whereas higher constitutive ESR gene expression (“frontloading” *sensu* ^8^) did not. Our results reveal a novel genomic mechanism of resilience to a variable environment, demonstrating that corals are capable of a more diverse molecular response to environmental stress than previously thought.

To explore expression patterns with respect to origin and transplant, we conducted a discriminant analysis of principal components for all genes represented by at least 10 unique transcript counts in more than 90% of samples. Corals from inshore populations exhibited greater plasticity in their genome-wide gene expression profiles than corals from offshore populations (Fig. 1). Inshore coral hosts transplanted to offshore reefs matched native offshore expression patterns more closely than offshore hosts transplanted to inshore reefs (t-test: t=2.94, df=11.7, P=0.01, Fig. 1A). *Symbiodinium* gene expression exhibited a similar trend, although it was not statistically significant (t-test: t=1.61, df=8.8, P=0.14, Fig. 1B).

**Figure 1.**
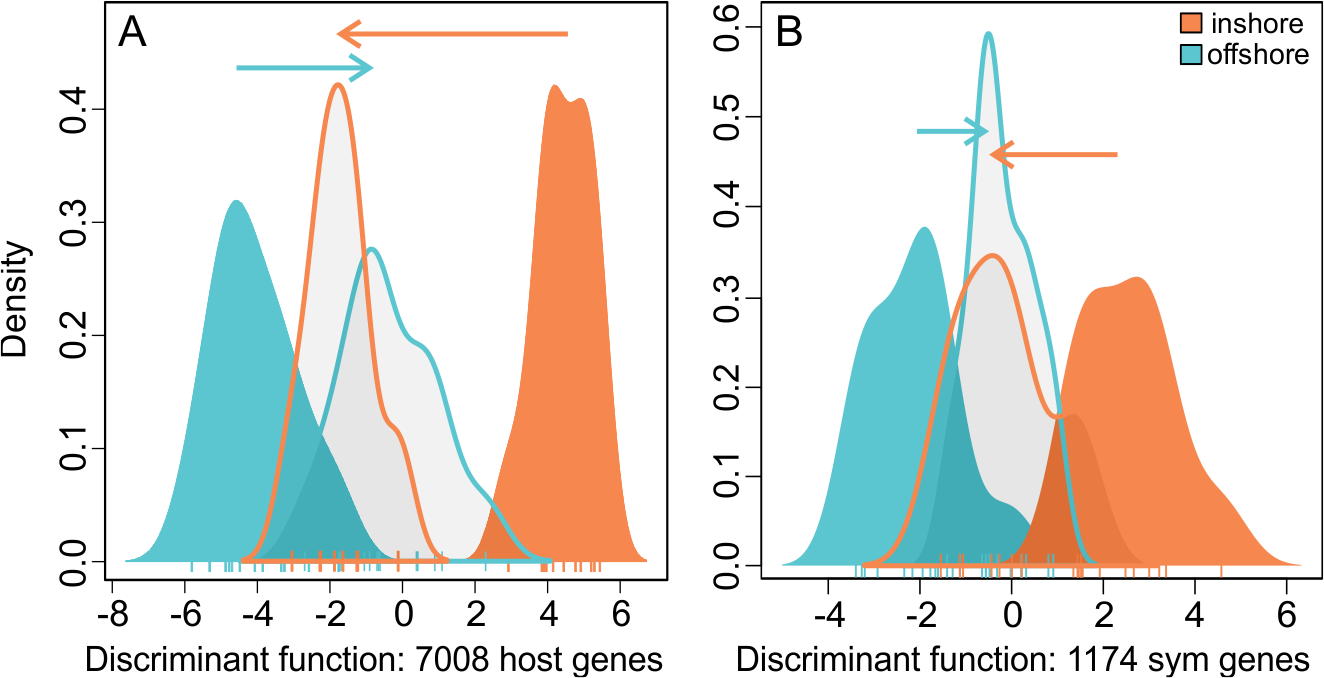
Population-level variation in genome-wide expression plasticity of coral hosts and *Symbiodinium.* (A) Discriminant analysis of principal components (DAPC) for coral host genes (n=7,008) for inshore and offshore corals when transplanted to their native reef (solid distributions) and non-native reef (transparent distributions) sites. (B) DAPC for *Symbiodinium* genes (n=1,174). Arrows indicate change in mean expression by population origin in response to transplantation.

To investigate these differences in more detail and explore their relationship with additional quantitative traits, we conducted weighted gene coexpression network analysis (WGCNA^7^) on the host and symbiont datasets. The 7,008 coral host genes were assigned to fourteen co-expression modules, four of which showed significant correlations with site of transplantation and the density of symbiont cells (Fig. 2A,B; n = 2,814 genes total). Eigengene expression (the first principle component of the expression matrix) of these modules again indicated that inshore corals exhibit more plastic expression than offshore corals (Fig. 3A,B) achieving a closer match to the expression profile of the native coral population upon transplantation (Fig. 3A,B).

**Figure 2.**
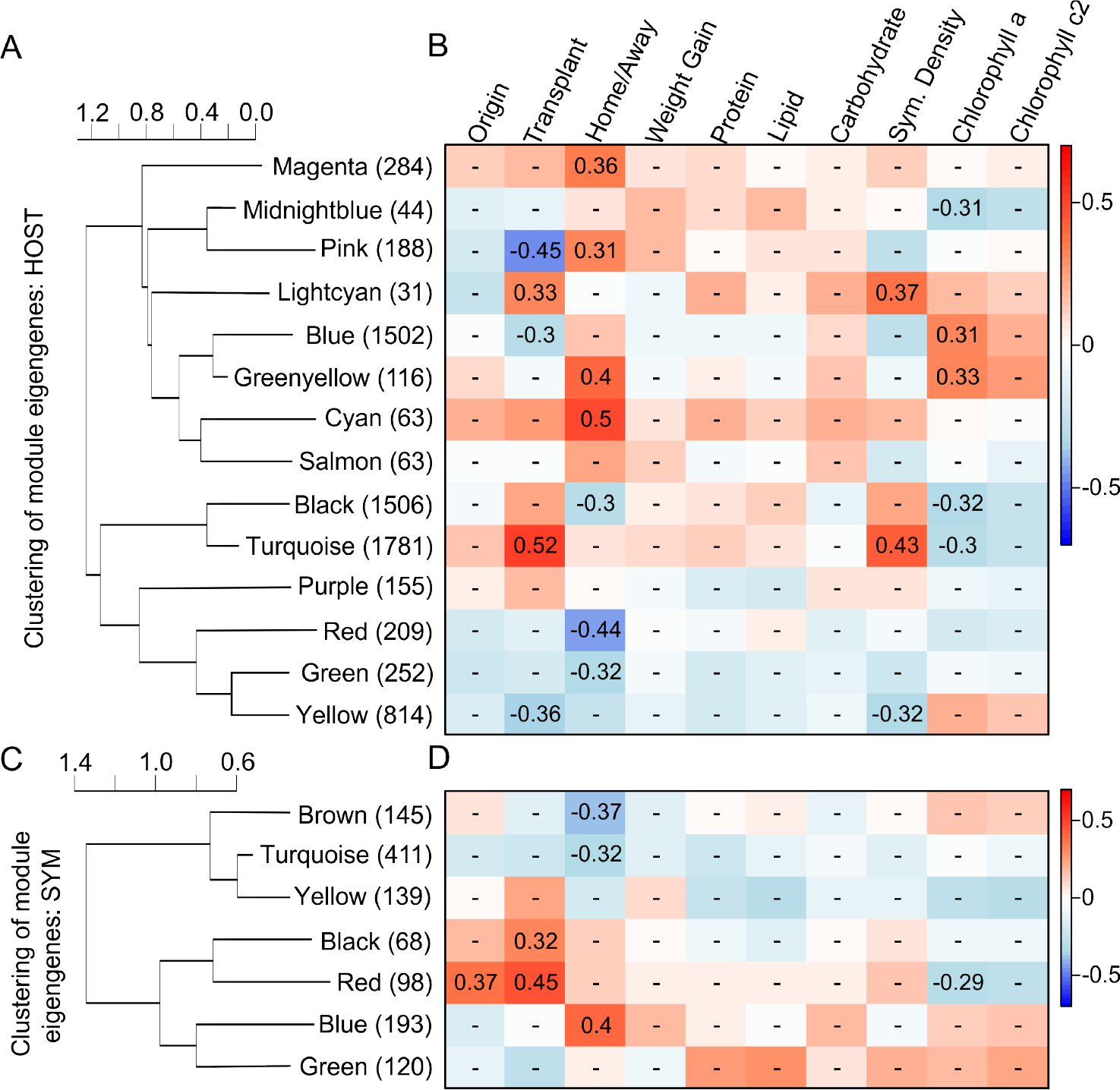
Hierarchical clustering dendrogram of module eigengenes (the first principal component of a module, representative of overall expression profile for genes within that module) and heatmaps of module-trait correlations for the coral host (A, B) and *Symbiodinium* (C, D). On panels A and C, the number of genes in each module is indicated in parentheses. On panels B and D, red indicates a positive correlation, blue a negative correlation, and Pearson’s *R* for significant correlations (P<0.05) are reported.

Functional enrichment analysis of the plastic host modules showing the strongest positive (Turquoise module) and negative (Yellow module) correlations with symbiont density indicated differential regulation of the environmental stress response (ESR) among populations. The top ‘biological process’ gene ontology (GO) enrichment for the Turquoise module was G0:0033554, ‘cellular response to stress’, and the top term for the Yellow module was GO:0042254, ‘ribosome biogenesis’, (Table S1). Taken together, these results demonstrate that corals transplanted to inshore reefs up-regulated cellular stress response genes, including molecular chaperones such as heat shock proteins, while down-regulating ribosome biogenesis, a hallmark of the ESR^9^.

**Figure 3.**
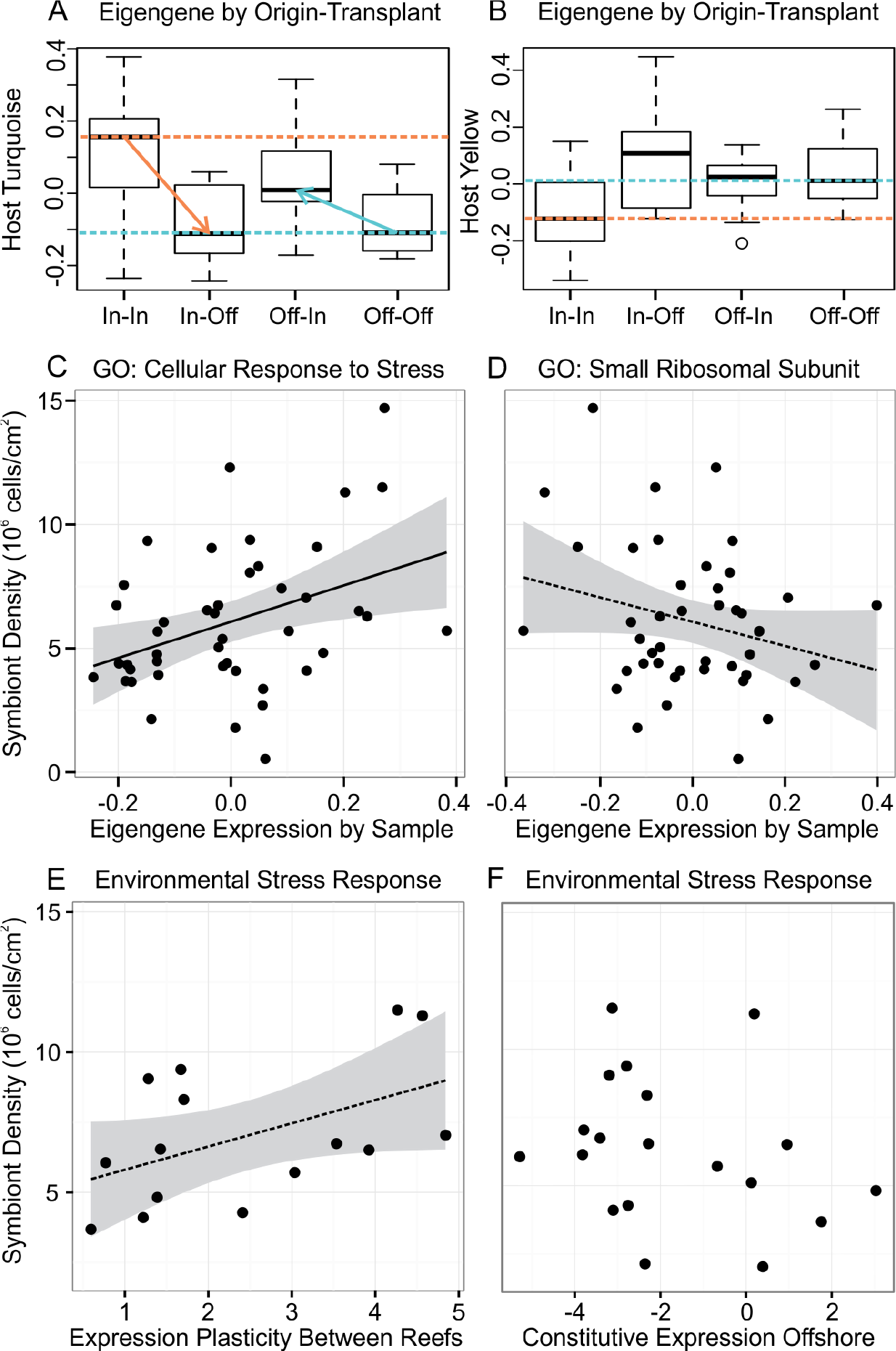
(A,B) Inshore coral hosts show greater gene expression plasticity of major symbiont-correlated modules than offshore corals. Boxplots show the median and interquartile range of module eigengenes (the first principle component of the expression matrix) with respect to site of origin and transplant environment for (A) host turquoise and (B) yellow modules. Colored dashed lines indicate origin-specific medians. (C, D) Corals with the strongest environmental stress response (ESR) also maintained the highest symbiont densities. Regression of symbiont densities on eigengenes for significantly enriched (P<0.001) gene ontology terms (C) ‘cellular response to stress’ (n=40 genes, host turquoise module, R: 0.39, 95%CI [0.10, 0.61], P<0.01) and (D) ‘small ribosomal subunit’ (n=17 genes, host yellow module, R: -0.26, 95%CI [-0.51, 0.04], P=0.09). (E) Coral genotypes with greater ESR expression plasticity maintained higher symbiont densities when transplanted to inshore reefs (R: 0.49, 95%CI [-0.03, 0.80], P=0.06). (F) Constitutive ESR expression level among genotypes was unrelated to maintenance of symbiont densities (R: -0.25, 95%CI [-0.64, 0.23], P=0.29).

Maintenance of homeostasis in the face of environmental variability is a physiological challenge faced by all organisms, but sessile animals, such as reef-building corals, are particularly susceptible since they cannot change habitats to escape stressors^9^. Inshore reefs in the Florida Keys are more variable than offshore reefs in multiple environmental parameters^10^, including temperature, and inshore corals exhibit elevated thermal tolerance^3,11^. In the present study, thermal stress was likely responsible for up-regulation of the ESR at inshore reefs, as samples were collected at the end of the 2012 summer^4^. In addition, coral bleaching, the stress-induced functional loss of the endosymbionts which commonly results from elevated temperature^12^, was observed at the time of sample collection, but only in offshore origin corals transplanted to inshore reefs (Fig. S2). Further supporting the conjecture that ESR regulation in the coral host is a direct response to elevated summer temperatures is the strong correlation between ESR expression and symbiont density: corals with the strongest ESR maintained the highest symbiont densities (Fig. 3C,D). Notably, corals with the greatest change in ESR expression across environments maintained the highest symbiont densities (Fig. 3E), while corals with higher constitutive expression of ESR genes did not (Fig. 3F). This indicates that differential thermal tolerance among populations was due to varying capacity for gene expression plasticity rather than elevated pre-emptive expression (“frontloading”) of stress-response genes, *sensu* ^8^. Furthermore, these results show that variation in bleaching susceptibility of the holobiont (the combination of host and symbiont) is explained by the ability of the coral host to mitigate intracellular damage resulting from environmental stress, which is in contrast to many other coral species where variation in bleaching susceptibility is largely attributable to differences in symbiont genotype (e.g.^13,14^).

In the *Symbiodinium*, 1,174 genes were assigned to seven co-expression modules, none of which showed correlations with symbiont density, though the red module was correlated with chlorophyll *a* content (Fig. 2C,D). Expression in two modules was correlated with site of transplantation, and expression plasticity in the red module tended to differ among populations, with inshore symbionts showing greater change than offshore symbionts. This module comprised few genes (n=98) with the most significantly enriched GO term being GO:0009521, ‘photosystem’ (Table S1). The genes in the module annotated with this GO term included components of the peripheral light-harvesting complex (LHC), such as fucoxanthin-chlorophyll *a/c* binding proteins. A decrease in peripheral LHCs limits the risk of photodamage to D1 reaction center proteins and has been proposed as a photoprotection mechanism of *in hospite Symbiodinium* in response to stress^15^. If *Symbiodinium* self-protection was the reason for elevated thermal tolerance of inshore-origin corals, inshore origin symbionts should exhibit down-regulation of these photosystem genes at inshore reefs where bleaching was observed (Fig. S2). However, the opposite pattern was evident: native inshore-origin *Symbiodinium* exhibited elevated expression in comparison to offshore-origin transplants (Fig. S3). This suggests that symbiont performance under thermal stress in inshore corals is maintained not due to the symbionts’ own stress protection mechanisms but through plasticity of host ESR genes.

Increased plasticity is predicted to evolve in a population if reaction norms vary across genotypes and the slope of the reaction norm is positively correlated with fitness^16,17^. Given that expression of stress response genes is energetically costly^18^, a tradeoff is expected where enhanced plasticity would be beneficial in the variable environment but detrimental in the stable environment. We estimated the costs of stress response expression plasticity using the selection gradient method^19,20^, by regressing relative fitness (weight gain) in the home environment against mean expression at home and the magnitude of expression change upon transplantation (plasticity). The partial regression coefficient for the plasticity term provides an estimate of cost while controlling for trait’s mean. A positive value indicates selection for plasticity, while a negative value indicates selection against plasticity^21^. The estimated plasticity coefficient for expression of cellular stress response genes (n=40 genes) was positive for inshore corals (0.11) and negative for offshore corals (-0.02), suggesting divergent selection acting among populations consistent with environmental variation among reef sites. However, since neither coefficient was significantly different from zero, additional work is needed to confirm this hypothesis.

Given the difference in thermal regimes between inshore and offshore reefs in the Lower Florida Keys it is expectable that inshore corals have adapted and/or acclimated to their native reef environment^4^ and exhibit elevated thermotolerance^3,11^. The surprising result of the present study is that these higher-order phenotypic responses may be explained by a differential capacity for gene expression plasticity. The population-level difference in gene expression plasticity described here is also in contrast to another recently reported mechanism by which corals may be adapting to temperature variation. Barshis *et al.* (2013) found that corals from more thermally variable pools exhibited constitutive up-regulation of ESR transcripts, which the authors termed “frontloading”. However, while ESR plasticity is associated with higher bleaching resistance (maintenance of *Symbiodinium* densities, Fig. 3E), we find no relationship between bleaching resistance and baseline ESR expression levels as observed in the same genotypes at the stable offshore reef environment; the trend is in fact negative (Fig. 3F). The difference between bleaching-resistance strategies (plasticity versus “frontloading”) may be due to the frequency at which coral populations are exposed to thermal stress events. The dominant cycle of temperature fluctuations in the Florida Keys occurs on an annual scale^4^, while the corals studied by Barshis *et al.* (2013) experience dominant fluctuations on a daily basis, during tidal cycles. Theory predicts that constitutive expression of an adaptive phenotype will be favored over plasticity if the environment fluctuates more rapidly than the typical response time^16^. The constitutive up-regulation of ESR genes by corals in tidal pools^8^ suggests that these populations integrate over the periodicity of stress events, analogous to a constant stress environment. In contrast, the variable expression of ESR genes in inshore coral populations observed here suggests that these corals have adopted an alternate solution, employing adaptive plasticity to cope with annual cycles of temperature variation in the Florida Keys.

Taken together, our results show that inshore corals from a more variable thermal environment developed an ability to more dynamically regulate expression of environmental stress response genes, which is associated with maintenance of *Symbiodinium* densities following thermal stress. Understanding the capacity of coral populations to adapt or acclimatize to local thermal stress is paramount for predicting coral responses to future climate change. Plasticity may accelerate evolution by facilitating mutational and genetic variance^22^ as well as by allowing coral populations to exploit novel emerging conditions^23^. Future work should aim to investigate how different strategies of constitutive “frontloading” and expression plasticity affect the capacity of coral populations to adapt to changing climates in the long term.

## METHODS

### Sample collection and processing

The transplantation experiment as been fully described in ^4^. Briefly, fifteen genotypes (individual colonies) of *Porites astreoides* from an inshore and an offshore reef in the Lower Florida Keys, USA, were fragmented and outplanted at native and foreign sites (Fig. S1) under Florida Keys National Marine Sanctuary permit #2011-115. Following one year of transplantation coral growth rates, energetic stores (total protein, lipid and carbohydrate content), symbiont densities and chlorophyll content were measured for each genotype. Immediately upon field collection, 1-cm^2^ tissue samples were taken from each coral fragment and preserved in RNALater (Ambion, Life Technologies) on ice. Samples were stored at -80°C until processing. Total RNA was extracted using RNAqueous kit (Ambion, Life Technologies), with minor modifcations. Briefly, samples homogenized in lysis buffer were kept on ice for one hour with occasional vortexing to increase RNA yields, which was followed by centrifugation for 2 minutes at 16100 rcf to precipitate skeleton fragments and other insoluble debris; 700 μl of the supernatant was used for RNA purification. At the final elution step, the same 25 μl of elution buffer was passed twice through the spin column to maximize the concentration of eluted RNA. Samples were DNAse treated as in ^24^ One μg of total RNA per sample was used for tag-based RNA-seq, or TagSeq ^5^, with modifications for sequencing on the Illumina platform. TagSeq was recently demonstrated to generate more accurate estimates of protein-coding transcript abundances than standard RNA-seq, at a fraction of the cost ^6^.

### Bioinformatic analysis

A total of 45 libraries prepared from each biological sample were sequenced on the Illumina HiSeq 2500 at UT Austin’s Genome Sequencing and Analysis Facility. Though 60 samples were originally outplanted, nine were lost due to hurricane damage ^4^ and six more were discarded during sample preparation for poor RNA or cDNA quality. Resulting sample sizes per origin to transplant group were inshore to inshore (n=11), inshore to offshore (n=9), offshore to inshore (n=13) and offshore to offshore (n=12). Overall, 527.9 million raw reads were generated, with individual counts ranging from 3.2 to 26.3 million per sample (median = 11.1 million reads, NCBI SRA:NNNN). A custom perl script was used to discard reads sharing the same sequence of the read and degenerate adaptor (PCR duplicates) and trim the leader sequence from remaining reads. The *fastx_toolkit* (http://hannonlab.cshl.edu/fastx_toolkit) was then used to trim reads after a homopolymer run of ‘A’ ≥ 8 bases was encountered, retain reads with minimum sequence length of 20 bases, and quality filter, requiring PHRED of at least 20 over 90% of the read. 0.3 to 2.1 million reads per sample (median = 0.9 million reads) remained after quality filtering. The *Porites astreoides* transcriptome ^25^ was concatenated to a *Symbiodinium* Clade A reference ^26^, as *P. astreoides* host A4/A4a-type symbionts in the Florida Keys ^3,27^. Filtered reads were mapped to this combined reference transcriptome with *Bowtie2* ^28^, using the ‒sensitive-local flag. Read counts were assembled by isogroup (i.e. groups of sequences putatively originating from the same gene, or with sufficiently high sequence similarity to justify the assumption that they serve the same function) using a custom perl script. Reads mapping to multiple isogroups were discarded. This count file was split into host-specific and symbiont-specific isogroup files for subsequent analyses. In total, 137,542 to 769,503 unique reads per sample (median=354,184 reads) mapped to 26,000 host isogroups and 7,393 to 66,623 unique reads per sample (median=26,760 reads) mapped to 21,257 symbiont isogroups.

### Differential expression, co-expression network and functional enrichment analyses

Analyses were carried out in the R statistical environment ^29^. Low expression genes (those with less than 10 counts in more than 90% of samples) were removed from the dataset, leaving 7,008 and 1,174 highly expressed genes in the host and symbiont datasets, respectively. Gene counts in both the host and symbiont datasets were normalized and log-transformed using a regularized log transform with the command *rlog*() in DESeq2 ^30^ for subsequent analyses.

A discriminant analysis of principal components (DAPC) was used to compare expression of all 7,008 highly expressed genes for host corals, and 1,174 genes for symbionts using the *adegenet* package ^31,32^. A discriminant function was built by defining native transplants as groups (inshore corals transplanted to inshore reefs, and offshore corals transplanted to offshore reefs). Group memberships were then predicted for the transplant samples based on the DAPC scores for the native populations. The difference between native and non-native expression was calculated within individual genotypes as a metric of expression plasticity and an unequal variances t-test was used to compare population means for hosts and symbionts.

WGCNA analysis was carried out following tutorials for undirected WGCNA ^7,33,34^ The analysis is blind to experimental design and involves four steps: (1) Pearson correlations for all gene pairs across all samples are computed to construct a similarity matrix of gene expression, retaining the sign of the expression change (“signed networks”); (2) Expression correlations are transformed into connection strengths (connectivities) through a power adjacency function, using a soft thresholding power of 6 for the host and 10 for the symbiont, based on the scale-free topology fit index (Fig. S4); (3) Hierarchical clustering of genes based on topological overlap (sharing of network neighborhood) is performed to identify groups of genes whose expression covaries across samples (network modules), retaining modules with at least 30 genes and merging highly similar modules (with module eigengenes correlated at *R*>0.85) (Fig. S5,6); and (4) External trait data is related to the expression of inferred modules (Fig. 2).

Functional enrichment analyses were conducted using the *GO_MWU* package in R (https://github.com/z0on/GO_MWU) to identify over-representation of particular functional groups within modules of the host and symbiont datasets, based on Gene Ontology (GO) classification ^35^. For each GO term, the number of annotations assigned to genes within a module was compared to the number of annotations assigned to the rest of the dataset, to evaluate whether any ontologies were more highly represented within the module than expected by chance (Fisher’s exact test).

### Selection gradient estimation

We quantified the costs of stress response expression plasticity for inshore and offshore populations by relating fitness to mean expression of the 40 genes comprising the ‘cellular response to stress’ functional enrichment (Table S1) at native reefs and the difference in expression between environments following ^19,20^. Relative fitness was defined as the growth of an individual genotype at its native reef relative to the maximum possible weight gained by a coral within each environment ^4^. Mean expression level and the difference in expression in response to transplantation (i.e. expression plasticity) were calculated for individual genotypes using standardized DAPC scores for this gene subset.

### Code availability

The current protocol and bioinformatics scripts for TagSeq are maintained at https://github.com/z0on/tag-based_RNAseq. R scripts and input files for conducting the gene expression and statistical analyses described here can be found on Dryad: NNNNN. R scripts and example input files for the GO_MWU package are actively maintained at https://github.com/z0on/G0_MWU.

## AUTHOR CONTRIBUTIONS

CDK and MVM conceived and designed experiments. CDK performed experiments, analyzed data and wrote the first draft of the manuscript. Both authors contributed to revisions.

## ACKNOWLEDGMENTS

G Aglyamova completed library preparation of samples for Illumina sequencing. M Strader created the GIS map of the Florida Keys. Funding for this study was provided by NSF DDIG award DEB-1311220 to CDK and MVM.

## Supplementary Material

**Figure S1.**
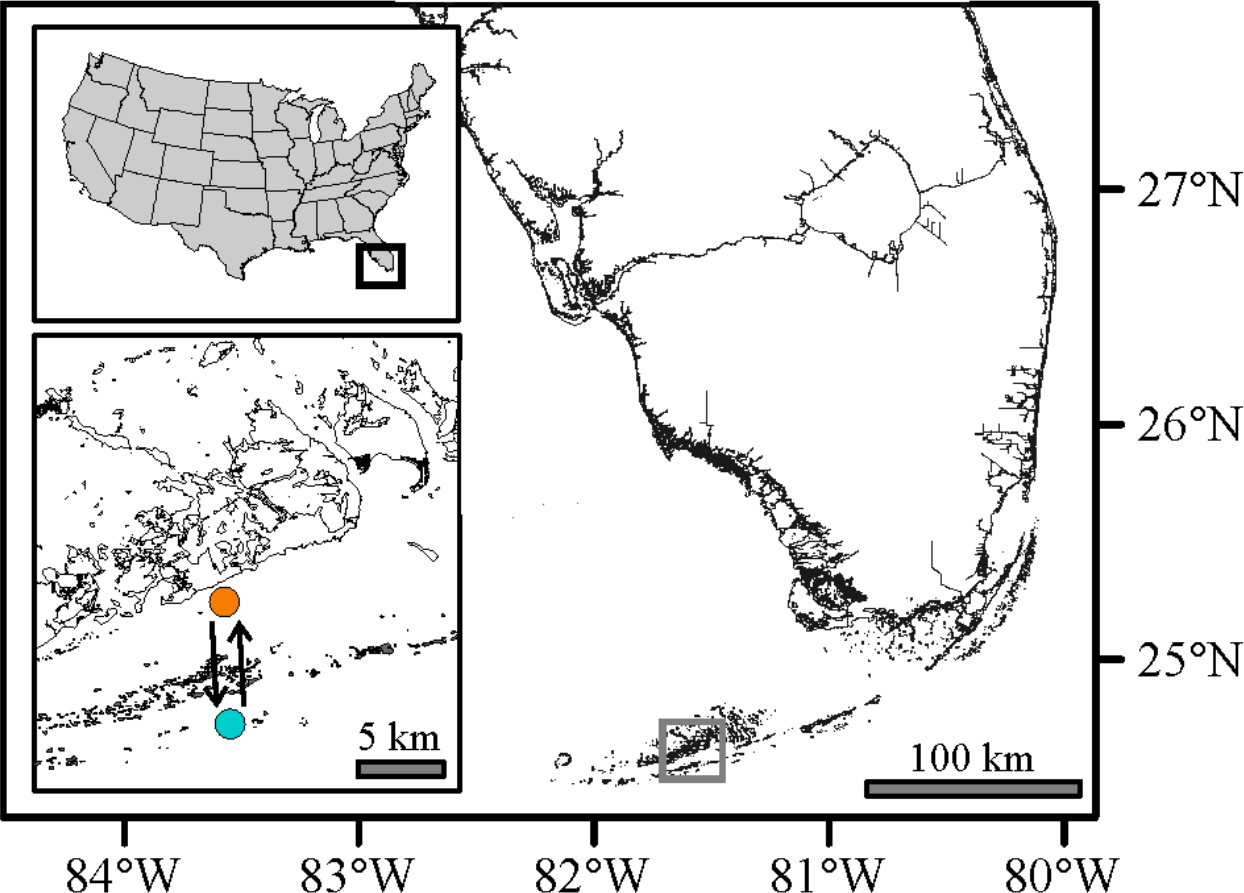
Map of the Florida Keys, USA, with inset showing reciprocal transplant sites for the Lower Keys.

**Figure S2.**
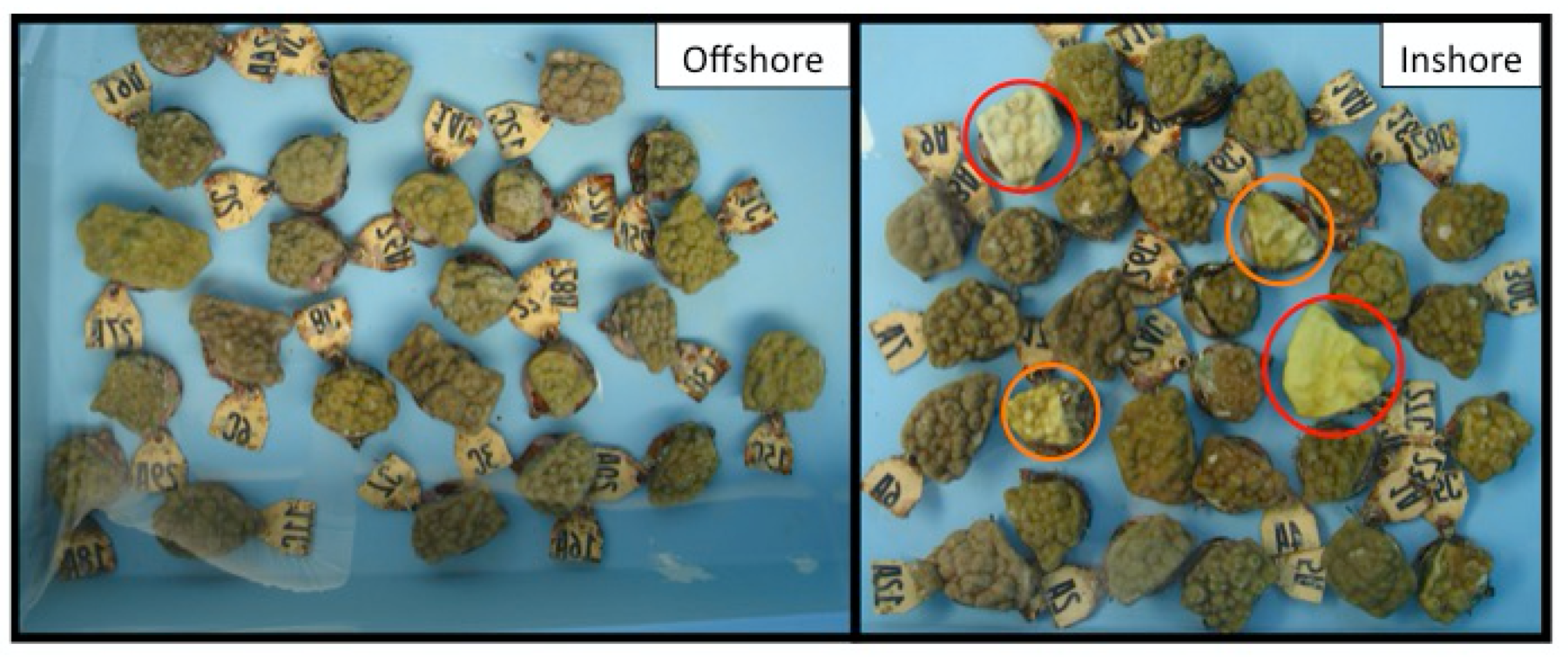
Photographs of coral fragments collected at the inshore and offshore site following one year of transplantation. Genotypes 1-15 are inshore origin; 16-30 are offshore origin. Four corals collected from the inshore site show signs of bleaching: genotypes 16 and 17 are pale (orange circles) while 21 and 27 are bleached (red circles).

**Figure S3.**
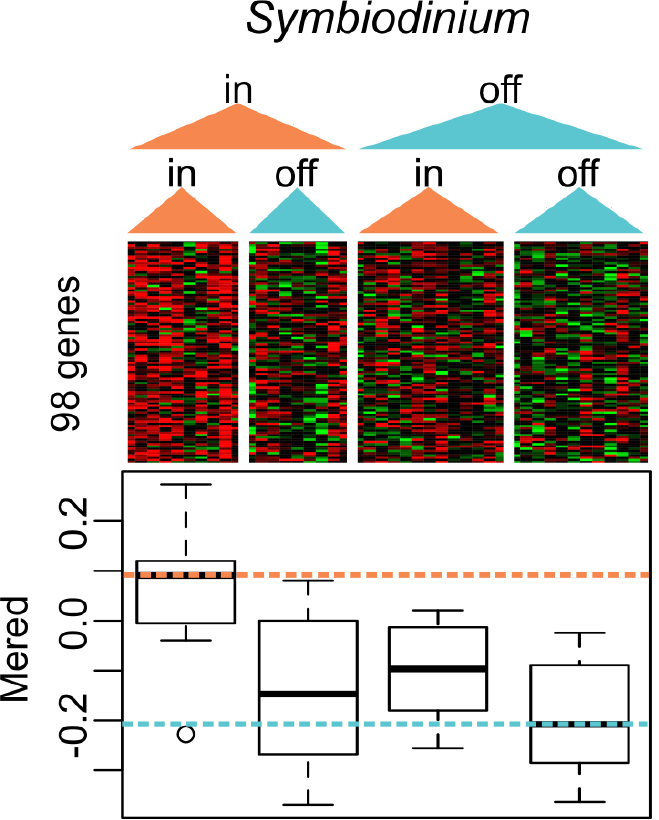
Heatmap of all genes assigned to the symbiont red module by individual samples. Boxplot shows the median and interquartile range of module eigengenes (the first principle component of the expression matrix) with respect to site of origin and transplant environment. Colored dashed lines indicate origin-specific means.

**Figure S4.**
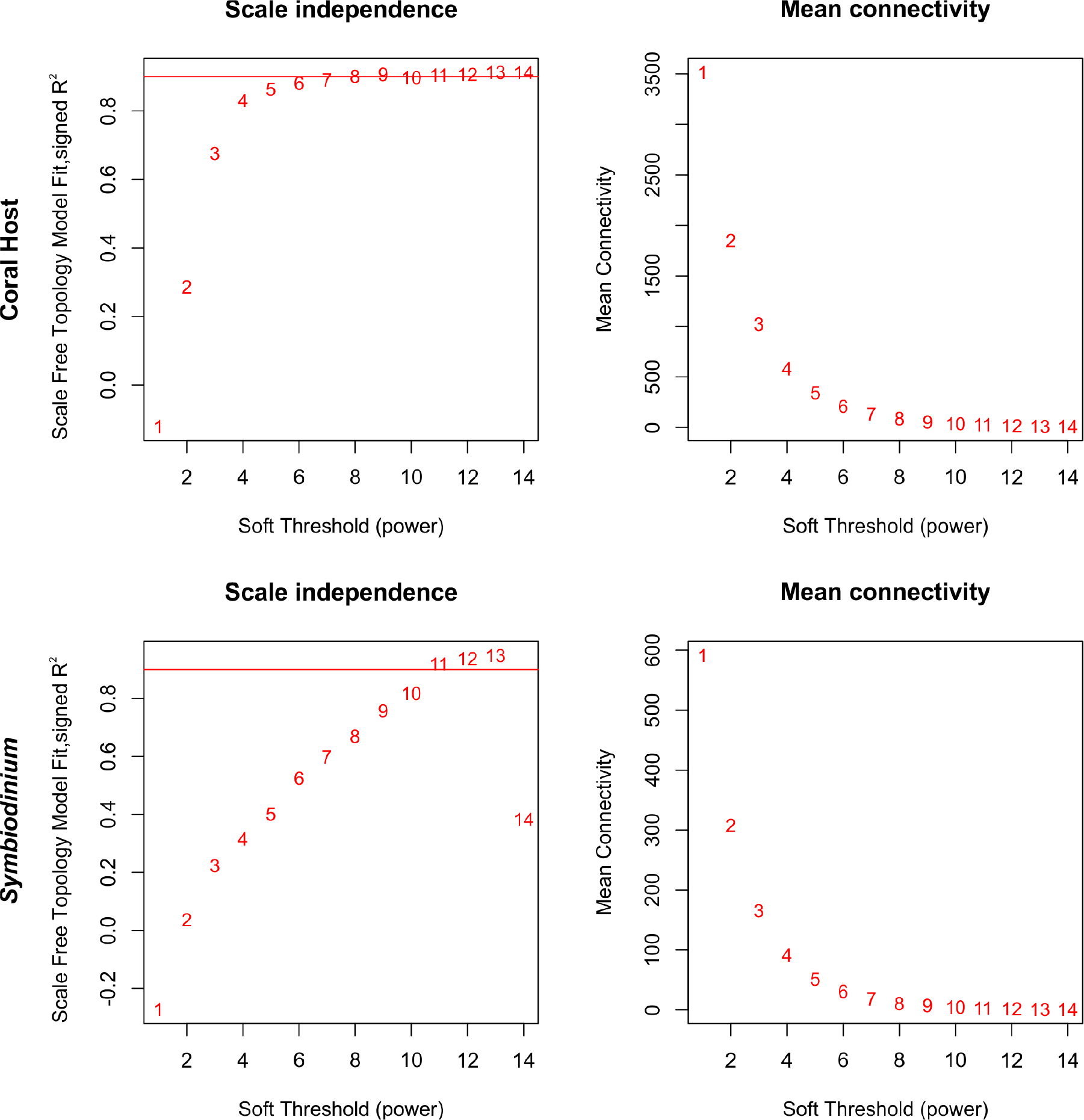
Analysis of network topology for select soft-thresholding powers. The scale-free fit index as a function of the soft-thresholding power is shown on the left, while mean connectivity as a function of the soft-thresholding power is on the right. We selected the power 6 for the host expression set and 10 for the symbiont expression set, the lowest values above 0.8 for which the scale-free topology fit index curve plateaus upon reaching a high value.

**Figure S5.**
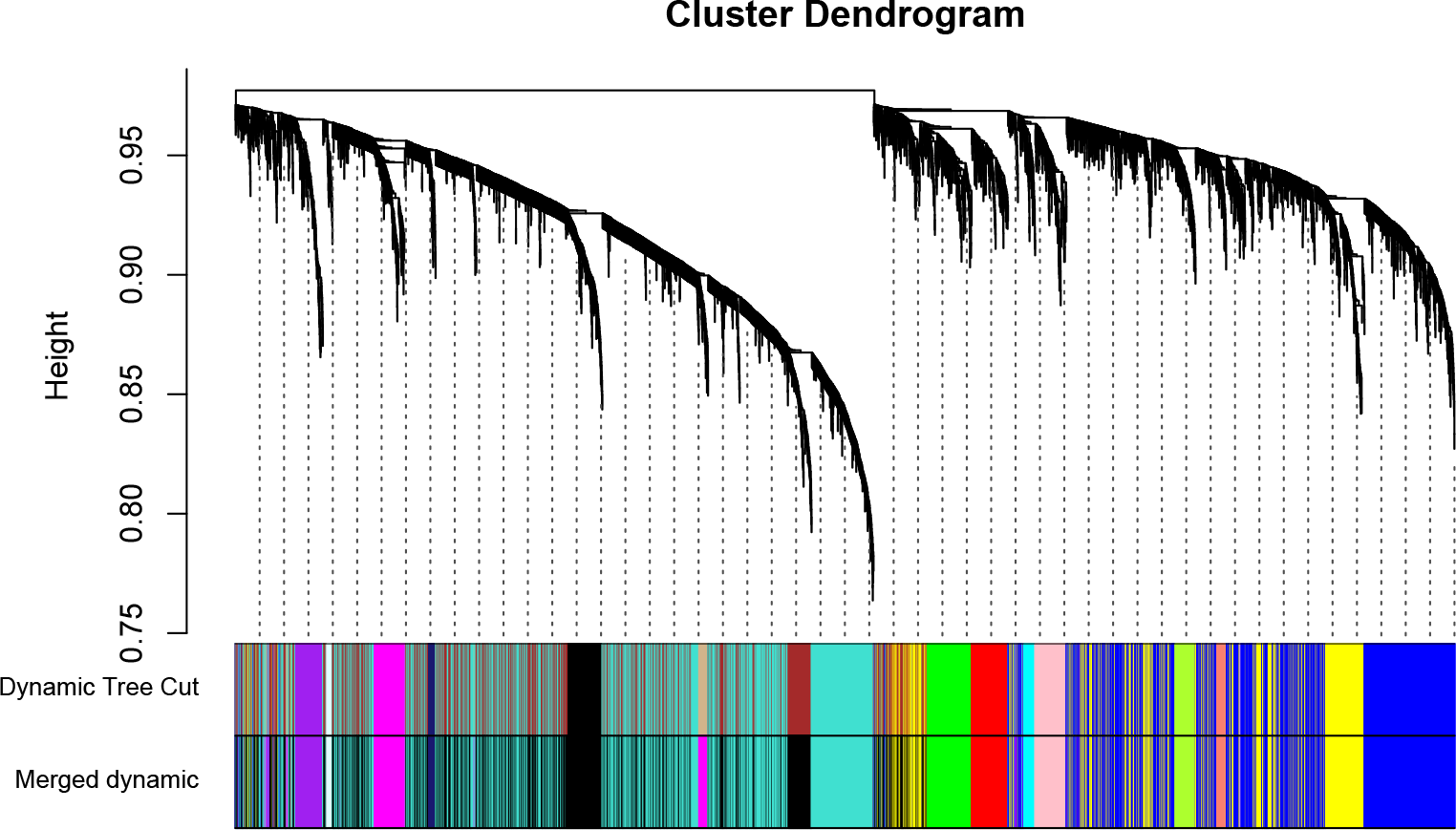
Clustering dendrogram of host expression set, with dissimilarities based on topological overlap shown with assigned module colors (Dynamic tree cut) and upon merging modules whose expression profiles were 85% similar (Merged dynamic).

**Figure S6.**
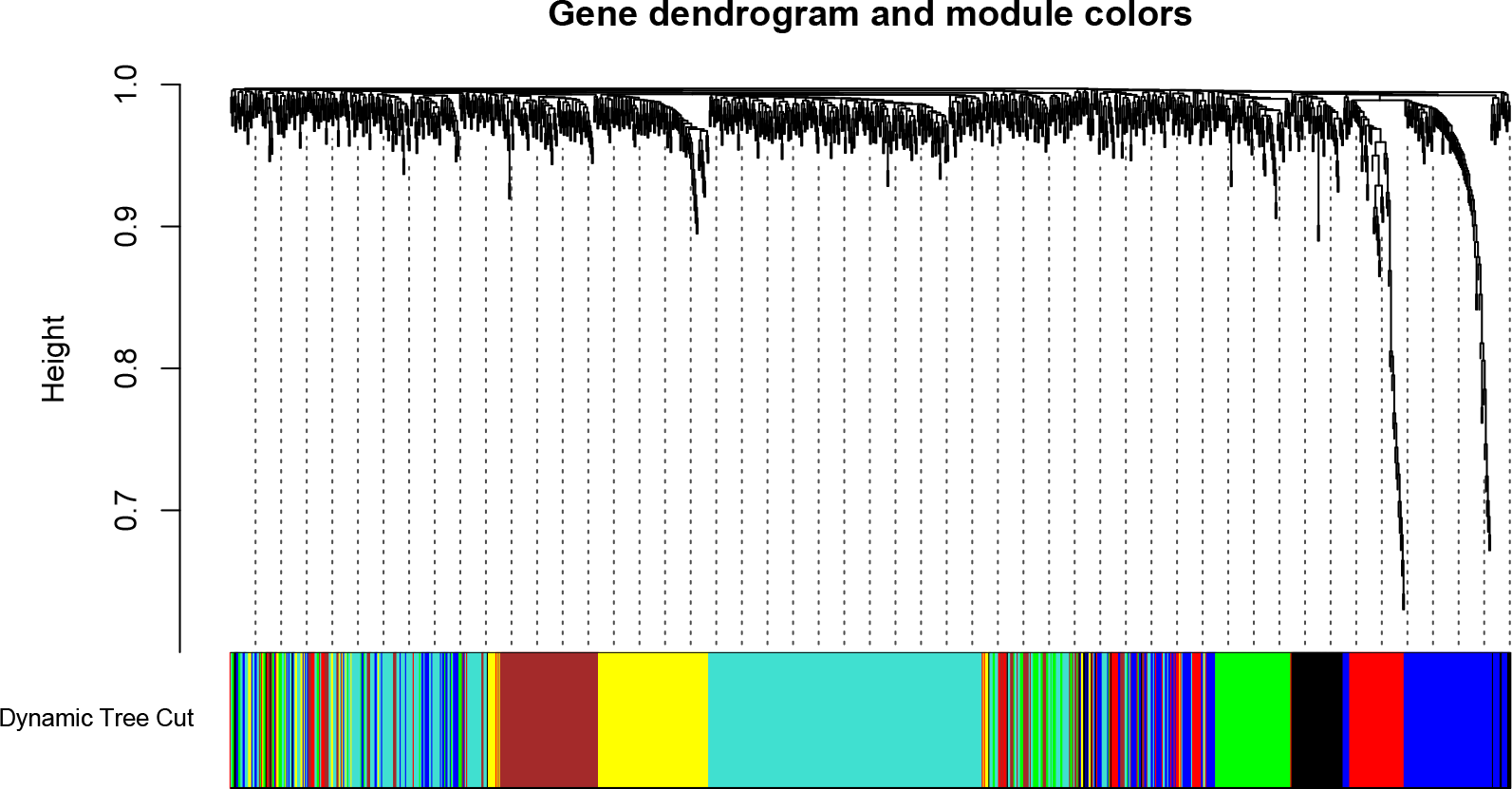
Clustering dendrogram of symbiont genes, with dissimilarities based on topological overlap shown with assigned module colors (Dynamic tree cut). No modules were similar enough to warrant merging.

**Table S1.**
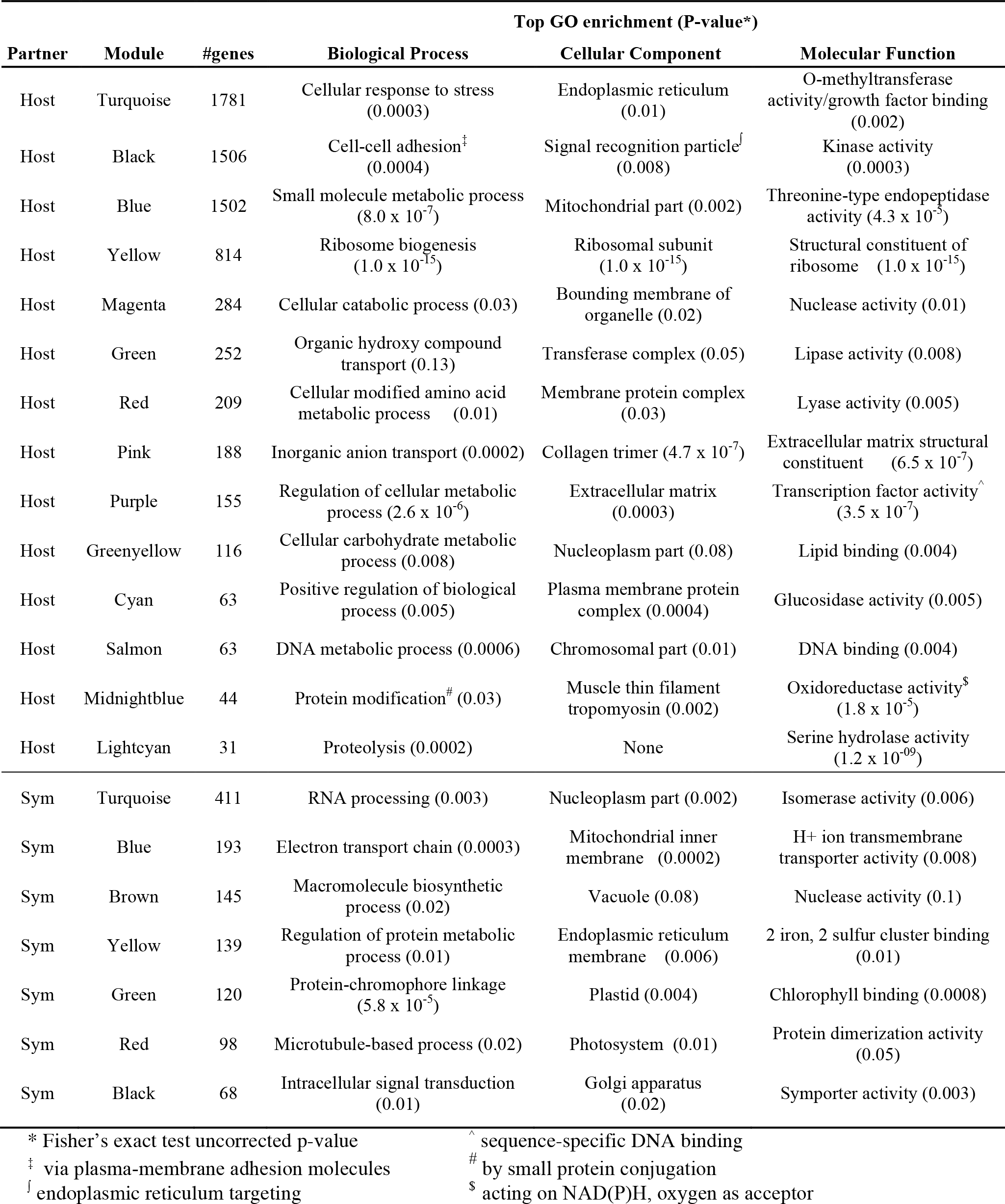
Top gene ontology (GO) terms resulting from functional enrichment analysis of significant co-expression modules for host and symbiont.

